# Novelty, category and orientation tuning for printed characters: a magnetoencephalography study with fast periodic visual stimulation

**DOI:** 10.64898/2025.11.29.691313

**Authors:** Ekaterina Kochetkova, Daria Kostanian, Olga Martynova, Olga Sysoeva

## Abstract

Letter recognition is assumed to involve several levels of analysis, including coarse tuning for category and novelty and more fine tuning, related to letter orientation. We employed an oddball fast periodic visual stimulation (FPVS) paradigm with magnetoencephalography (Elekta VectorView, 306 sensors) to study neural discrimination responses in the source space. Using contrasts between native letters and foreign letters, digits, or inverted native letters, we aimed to isolate the neural responses to visual novelty, category, and orientation during character analysis. The study was conducted with a cohort of 25 adults. The response topography demonstrated bilateral organization, including language-related brain regions as ventral occipitotemporal cortex, inferior parietal cortex and middle temporal areas. Comparing conditions, we revealed right lateralized parietal clusters, associated with novelty tuning, and left lateralized occipitotemporal clusters exhibiting higher activity for letters among digits discrimination, supporting the role of this area in letter processing. No distinct spatial patterns specific to orientation tuning were observed in comparison to novelty and category tuning. We proposed that expertise-dependent orientation-specific tuning mechanisms may operate within an embedded, spatially overlapping with coarse tuning neural framework, characterized by special spatiotemporal patterns.

## Introduction

Among all visual graphical characters, letters are the most significant to humans. As structural units of written words, letters provide the basis for written language. Letter recognition is associated with the special systems of the brain. According to plenty neuropsychological, electrophysiological, and neuroimaging studies, it is established that the visual processing of letters and words involve visual word form area, the brain region in the left inferior temporal cortex (Lochy et al., 2018; McCandliss et al., 2003; Schlaggar & McCandliss, 2007), but may be not limited to the single region, including system of interacting brain areas (McCandliss et al., 2003; Pollmann et al., 2003; Price & Devlin, 2011; Wang et al., 2021).

The distinction between letters and other characters, however, is mainly driven by the experience of interaction with them that is also related to the cultural background (Polk & Farah, 1998; Testolin et al., 2017). In particular, letters are almost indistinguishable from the other categories to infants or illiterate individuals (Dehaene & Cohen, 2007). Acquiring written language skills leads people to develop special systems of letter recognition. The first basic level of letter recognition is known as a coarse neural tuning for print. This type of tuning is well described in previous studies and is related to sensitivity to letters compared to other characters such as symbols, shapes, pseudofonts, and foreign language letters (Maurer et al., 2006). Coarse neural tuning for print seems to develop quite early, even before the actual acquisition of reading skills, and is most likely related to the visual experience of interacting with letters in the environment (Cantlon et al., 2011; Kostanian at al. 2025). However, in addition to coarse tuning for letters, it has been suggested that there is another level of tuning for letters related to more fine specific features of letters, such as their orientation (Tong et al., 2016; Xue et al., 2023). It starts to develop only at a sufficiently advanced level of letter expertise, when reading skills have already been acquired (Kostanian at al. 2025).

The brain systems behind the coarse neural tuning to letters and tuning for letters orientation can be studied using magnetoencephalography (MEG). The main advantage of using MEG is its high spatial resolution without sacrificing temporal resolution. Oddball fast periodic visual stimulation (FPVS), or “frequency-tagging” (Liu-Shuang et al., 2014; Rossion et al., 2015) is a powerful new approach in MEG and EEG studies of visual stimuli recognition. It allows investigating discriminability between different stimuli categories in a short experimental procedure that does not require any challenging behavioral task. This approach is based on steady-state visual evoked potentials: when periodic visual stimulation at a specific frequency elicits a brain response at exactly the same frequency and its harmonics (Norcia et al., 2015). The oddball variation of FPVS approach implies that a sequence of base stimuli (presented at a certain frequency, e.g., 6 Hz) contain a rare deviant stimuli presented at a lower frequency (e.g., every fifth stimulus, 6 Hz/5 = 1.2 Hz). In such cases if deviant stimuli are identified as different from base ones, the neurophysiological response would also contain peaks not only at frequency of a base stimulation, but also at frequency of a deviant (lower) stimulation and its harmonics. This discrimination response at the frequency of deviant stimulation represents neurophysiological measures of discriminability (Norcia et al., 2015).

While many previous studies employed FPVS approach to study word (De Rosa et al., 2024; Hauk et al., 2025; Lochy et al., 2015, 2024; Lutz et al., 2024; Pescuma et al., 2022; van de Walle de Ghelcke et al., 2020; Wang et al., 2021) and letter processing (Barzegaran & Norcia, 2020; Crollen et al., 2025; Lochy & Schiltz, 2019; Kostanian et al., 2025), several unresolved questions remain. First, no study investigated comparative localization of brain systems, responsible for different levels of tuning for letters. To examine this question, we include not only conditions with unfamiliar graphical characters (foreign letters) which would allow us to assess course tuning effects, but also conditions with inverted letters of native alphabet to assess tuning for orientation. Second, it remains unclear whether letter-selective responses reflect coarse categorical processing itself, or whether they may be category selective for other familiar but not linguistic symbols (i.e. digits). Therefore, we consider the comparisons of the response for letter and digit stimuli as categorical contrast. Third and lastly, we consider the question of how to interpret VFT results by using switched categories of standards and deviant stimuli. If the response to deviant stimuli reflects only mismatch detection, replacing standard and deviant stimuli will not produce significant differences. Otherwise, we may suggest that the discrimination of deviant stimuli may depend on the background composed by standards and should be taken into account in future research. Currently, only two FPVS studies (Lochy & Schiltz, 2019; Lutz et al., 2024) have considered conditions with switched categories of standards and deviant stimuli and have shown that switching the standard and deviant categories can lead to controversial results. For this reason, it is important that each of our conditions has a matching opposite condition in which the standard and deviant stimulus categories are swapped.

Based on the above research questions, we implicated six experimental conditions of oddball FPVS sequences presented to native Russian adult speakers. We systematically varied three factors: novelty (familiar vs non-familiar, i.e. letter of native, Russian alphabet vs with letters of Georgian alphabet), category of stimuli (letter vs digit) and orientation (upright vs inverted).

## Methods

### Participants

The study included 28 adult native Russian speaker volunteers (18 females, all 28 were right-handed) with normal or corrected-to-normal vision. The sample size was calculated using the G*power 3.1.3 software (Heinrich-Heine-University, Düsseldorf, Germany) with a statistical power of 80%, and a significance level of 0.05. To assess effects using a paired T-test (anticipated effect size = 0.5, according to approximate medium effect size), minimum sample size should be 27 people. Three participants were excluded from analysis due to the lack of individual MRI data. Final analysis was therefore conducted on the remaining 25 participants (16 females, mean age = 27.0 ± 4.6).

All participants have given written informed consent. The study, approved by the Institute of Higher Nervous Activity and Neurophysiology’s ethics committee (June 30, 2023), followed the Helsinki Declaration’s ethical guidelines for human research. The study was conducted at the unique research facility “Center for Neurocognitive Research (MEG-Center)” of MSUPE.

### Stimuli and experimental procedure

Four categories of stimuli were used in the study: eight uppercase letters of the Cyrillic alphabet (А, Л, К, З, В, Р, Я, Ч); eight letters of the Georgian alphabet (ზ, ს, წ, დ, რ, ჰ, ც, ტ); eight digits (1, 2, 4, 5, 6, 7, 8, 9) and eight inverted Cyrillic letters (rotated by 180° upside-down). Perimetric complexity (the ratio of inked surface to the perimeter of this inked surface (Attneave & Arnoult, 1956)) was 0.496±0.126 for Cyrillic letters; 0.594±0.265 for Georgian letters and 0.255±0.092 for Digits. Stimuli were presented on a black background in white Times New Roman font at size 315 pt. Each letter was placed inside a star-shaped white frame (line width 6 pixels). The average letter size was 3°34’47’’×3°7’57’’ degrees of visual angle and the size of the star-shaped frame was 13°26’36“×13°26’36”. Images were presented at the center of the screen.

The stimulation sequence had the following structure: standard stimuli were presented at 6 Hz (six stimuli per second), and every fifth item was a stimulus of the deviant category presented at 1.2 Hz, such as BBBBDBBBBD. The stimuli were randomly chosen from the stimuli set, with no immediate repetition of the same stimulus. Each stimulation sequence started with 2 s of gradual stimulation fade in, 120 s of stimulation sequence, and 1 s gradual fade out. Each sequence included 720 stimuli: 72 repetitions of 8 base letters, and 18 repetitions of 8 deviant letters. Sequences were created and executed using PsychoPy software (v2022.2.5) (Peirce, 2007). To maintain a constant level of attention throughout the stimulation, participants performed a color-change detection task. The instruction was to press the space button at the moment when the color of a star-shaped frame that surrounds each letter changed. During the experiment, the frame turned from white to yellow for 0.5 s at random time points 14 times per sequence.

The name of the condition was coded based on three levels of stimuli analysis introduced in our study. In particular, a first letter corresponds with orientation analysis and can be either U(pright) or I(nverted), a second letter corresponds with novelty analysis and can be either F(amiliar) or N(on-familiar), third letter refers to the category analysis and can be either L(etter) or D(igit). Three conditions contain Cyrillic letters (Upright Familiar Letter, UFL) as a base stimuli category and non-native Georgian letters (Upright Non-familiar Letter, UNL), or digits (Upright Familiar Digit, UFD) or Cyrillic letters rotated by 180° clockwise (Inverted Familiar Letter, IFL) as deviant categories. Three switched conditions contain Georgian letters (UNL), digits (UFD) or inverted Cyrillic letters (IFL) as base stimuli and upright Cyrillic letters (UFL) as deviant stimuli. See a schematic representation of the experimental design in Fig. 1. In addition to general effects of novelty (first line, Fig. 1), category (second line, Fig. 1), and orientation (third line, Fig. 1) we compared the brain responses between conditions.

**Fig. 1.**
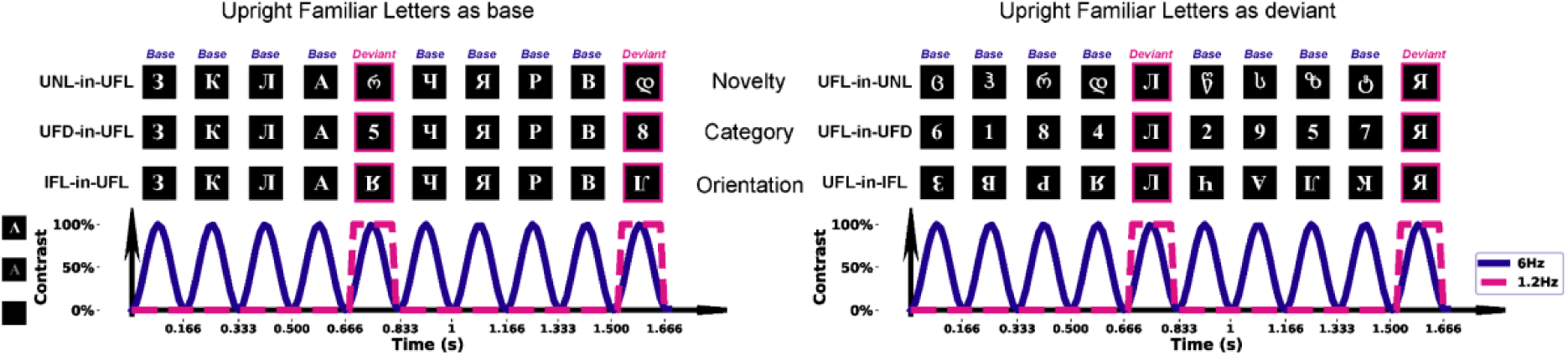
Schematic representation of the experimental paradigm. There were six conditions: Upright Non-familiar Letter in Upright Familiar Letters (UNL-in-UFL); Upright Familiar Digit in Upright Familiar Letters, (UFD-in-UFL); Inverted Familiar Letter in Upright Familiar Letters (IFL-in-UFL); Upright Familiar Letter in Upright Non-familiar Letters (UFL-in-UNL); Upright Familiar Letter in Upright Familiar Digit (UFL-in-UFD); Upright Familiar Letter in Inverted Familiar Letters (UFL-in-IFL). Each stimulation sequence lasted 120 s, during which stimuli were presented by sinusoidal contrast modulation at 6 Hz. Deviant stimuli appeared at 6/5 Hz (1.2 Hz) and highlighted in pink.

### MEG data acquisition and preprocessing

MEG data were collected at the Moscow Center for Neurocognitive Research using an Elekta VectorView 306 sensor (204 planar gradiometers and 102 magnetometers). MEG signals were recorded with built-in filters of 0.03–330 Hz and sampling rate of 1000 Hz. Noisy channels were identified via visual inspection. The signal was preprocessed with MaxFilter software (version 2.2.15) to reduce external noise using the temporal signal-space separation method (tSSS) and to realign data from different blocks to a standard head position. The following analyses were performed using MNE python software (Gramfort et al., 2013). Raw data was band-pass filtered between 0.1 and 100 Hz and notch filter at 50 Hz was applied.

### Frequency Domain Analysis in the Source Space

For the source space estimation we used individual MRI data pre-processed with Freesurfer V6.0.0 (Fischl, 2012). Anatomical T1-weighted magnetic resonance images were acquired during a separate session in MR scanner ‘Az-400’ (AO NPF ‘Az’) using 3D MPFIns sequence, TR = 1321 ms, TE = 11 ms, TI = 100 ms, 1 mm3 resolution, scanning time 11 min. The head model (3-layer boundary element model) was created using MNE-python. Source estimation was conducted using L2-minimum-norm estimation with a loose orientation constraint (regularization parameter based on a SNR value of 3), following the methodology similar to that applied in (Hauk et al., 2021).

For frequency domain analyses, Fast Fourier Transform (FFT) with frequency resolution 0.0083 Hz (1/120 s) was applied. From the acquired FFT spectrum we derived 10 segments centered at the frequency of interest and its nine higher harmonics, which included 21 frequency samples at each side of the center frequency, similarly to the methodology used in (Hauk et al., 2021). To estimate SNR, amplitude at each frequency bin was divided by the average amplitude of 20 surrounding bins (10 on each side and a gap of one frequency bin on each side of the target frequency to avoid spectral leakage) excluding the immediately adjacent and the extreme (min and max) bins (Liu-Shuang et al., 2014). The baseline correction of noise levels around each frequency of interest was applied by averaging the amplitude of ten neighbouring frequency-bins on each side of the center frequency and subtracting it from each frequency bin (Hauk et al., 2021). Finally, Z-scores were calculated for the sum of center frequency estimates and its first nine harmonics (from 1.2 Hz until 14.4 Hz for the frequency of deviant stimulation, harmonics associated with frequency of base stimulation (6 and 12 Hz) were excluded) according to methodology implemented in (Hauk et al., 2021). The number of harmonics were chosen according to the recommendations in (Retter et al., 2021), meaningful frequency range limitation and maximum number of harmonics for which we observed significant response for the deviant stimulation frequency (see Fig. S1 in Supplementary materials).

For the source space the significance of responses (sums of harmonics at frequency of base and deviant stimulation) at group level for each condition, was determined using cluster-based permutation testing. At the level of each condition we used a one-tailed t-test to define clusters at p-value threshold of 0.00001 and the significance of clusters was defined with permutations (n = 10000) using cluster-p-threshold of 0.05. Then analysis was continued according to the schema provided in Fig. 2. These comparisons of neurophysiological responses at the frequency of deviant stimulation in different conditions were also conducted using cluster-based permutation testing (two-tailed t-tests, with p-value threshold to form cluster of 0.001, significance of clusters was defined with permutations (n = 10000, cluster p-threshold of 0.05). For that purpose, we compared the summed Z-scored values at frequency of deviant stimulation (central 1.2 Hz frequency, summed up with its first nine harmonics, except 6 and 12 Hz) in pairs of conditions.

To investigate the difference of lateralization in conditions the laterality index (𝐿𝐼 = (𝐿𝐻 − 𝑅𝐻)/(𝐿𝐻 + 𝑅𝐻)) was calculated based on individual Z values for summed harmonics for frequency of deviant and base stimulation. Values for analysis were extracted for the selected regions of interest (ROI) that were shown in literature to be associated with reading and processing of linguistic information (De Rosa et al., 2024; Lochy et al., 2015; Pescuma et al., 2022; Pollack & Ashby, 2018; Wang et al., 2023) and was shown to reveal significant response in the majority of conditions in the current study, based on previous analysis in source space. The values for chosen ROI (according to Desikan-Killiany atlas embedded in MNE parcellation: ‘lateral occipital’, ‘fusiform’, ‘inferior parietal’, ‘inferior temporal’, ‘middle temporal’, ‘parahippocampal’, ‘lingual’, ‘transversetemporal’, ‘banksst’) were extracted using maximum value over the vertices values related to the certain parcel. An LI score above zero was interpreted as left lateralization, and a LI score below zero as right lateralization. T-test against zero for the mean laterality index value (across subjects) was used to define statistically significant lateralization of the response.

**Fig. 2.**
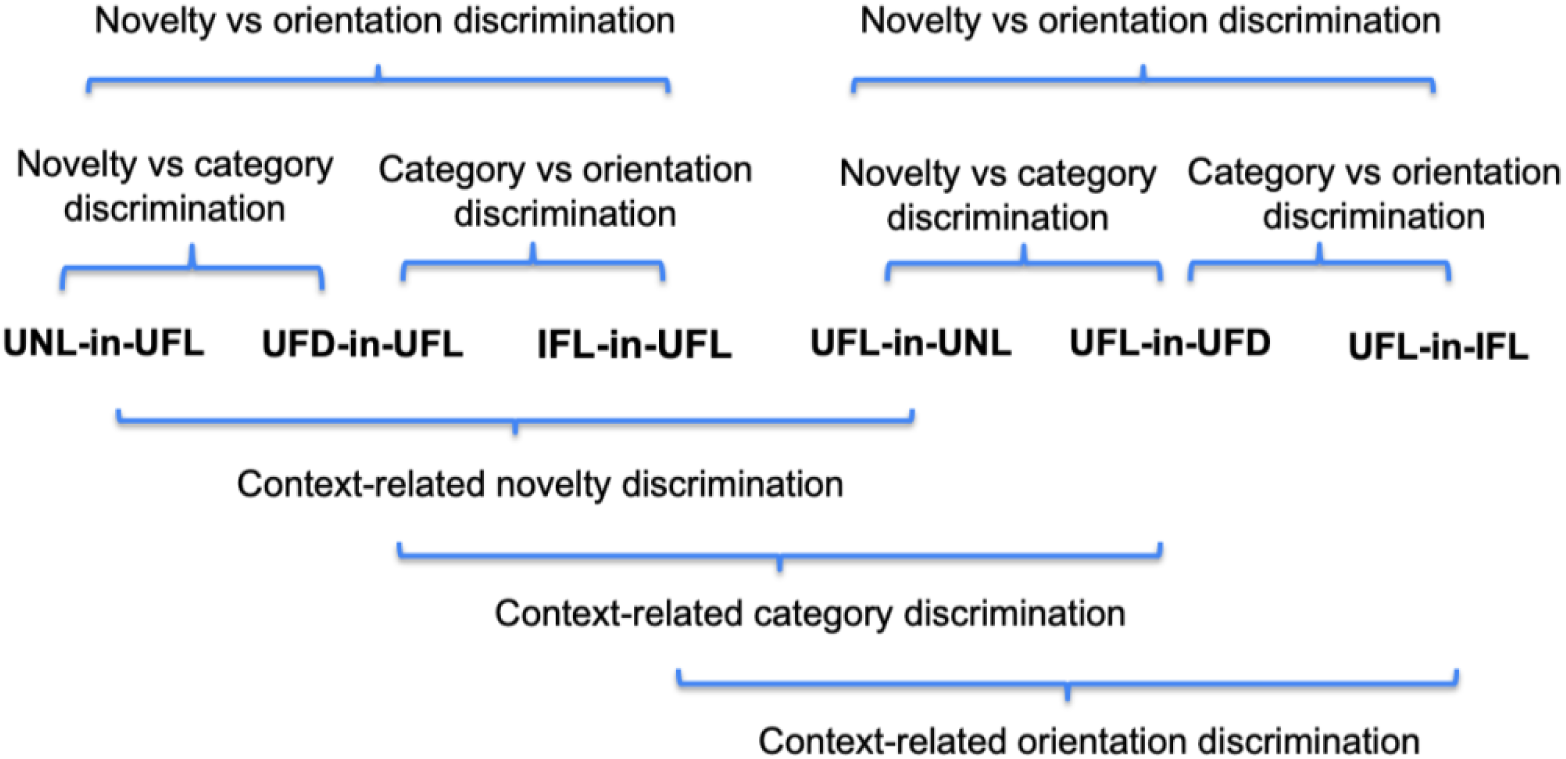
Schematic representation of analysis pipeline. Further analysis involves several comparisons between conditions: 1) comparisons of UNL-in-UFL and UFD-in-UFL, UFL-in-UNL and UFL-in-UFD to differentiate novelty effect from categorical discrimination, 2) comparisons of UFD-in-UFL and IFL-in-UFL, UFL-in-UFD and UFL-in-IFL, to analyze the difference between fine tuning for orientation and categorical discrimination, 3) comparisons of UNL-in-UFL and IFL-in-UFL, UFL-in-UNL and UFL-in-IFL to disentangle the difference between novelty effect and tuning for orientation, 4) comparisons between switched blocks as UNL-in-UFL vs UFL-in-UNL, UFD-in-UFL vs UFL-in-UFD, IFL-in-UFL vs UFL-in-IFL to test the context-dependent effects of novelty, category and orientation discrimination respectively.

## Results

### General overview of discrimination response and its lateralization

Significant discrimination responses in the source space for the frequency of deviant stimulation (1.2 Hz and its first nine harmonics, except multiplies of the base harmonics, i.e. 6 Hz and 12 Hz, see Methods for details) were found for all conditions in the lateral occipital, inferior parietal and inferior temporal regions, including ventral occipito-temporal cortex (vOTC) bilaterally (see Fig. 3).

Topographies of the response to the base stimuli (fundamental frequency of 6Hz and its harmonics) is given in Supplementary materials (Fig. S2). Since the response for base frequency of stimulation may represent the primary neural entrainment to the periodic visual input (Łabęcki et al., 2024) and the responses did not significantly differ between conditions, next we will mainly focus on the discrimination response (at the frequency of deviant stimuli).

Brain regions were considered as involved in the discrimination response when the significant clusters from the permutation cluster test showed over 50% overlap with parcellation labels. The regions involved in the discrimination response are summed up in Table 1. We can observe that more brain areas were activated for novelty condition compared to category and orientation conditions. The left precuneus demonstrated more prominent activation at orientation but not category contrast. At the same time the switched conditions were largely similar at least within the left hemisphere. Some areas were active in all conditions (i.e. fusiform area, inferior parietal and lateral occipital regions). It is noteworthy that the majority of research does not go further than this mostly descriptive analysis, while we persist in identifying statistically significant clusters that differentiate conditions.

**Fig. 3.**
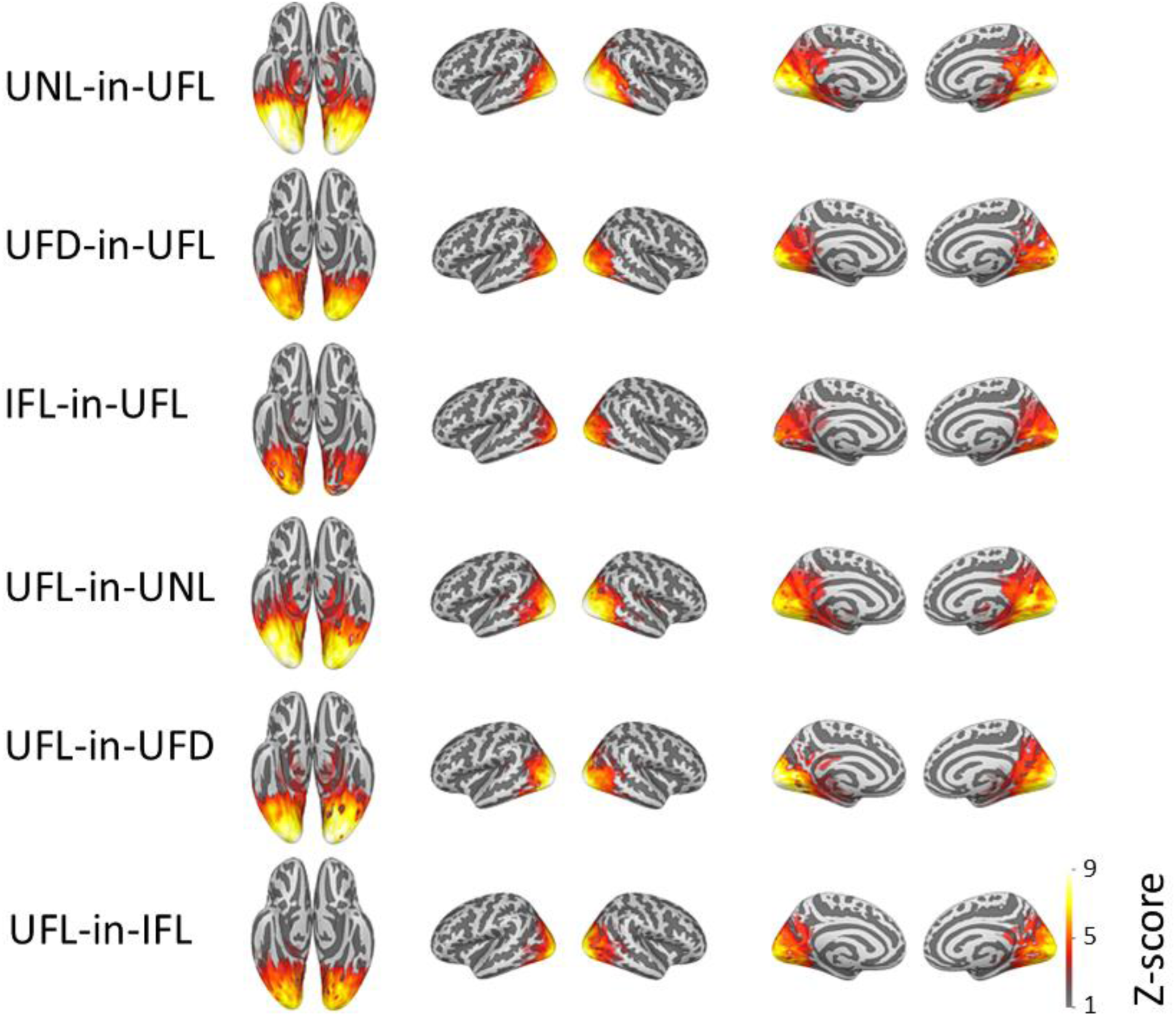
Source space analysis. Z-scores for the summed frequency of interest (1.2 Hz) and its nine harmonics for frequency of oddball stimulation at all experimental conditions: Upright Non-familiar Letter in Upright Familiar Letters (UNL-in-UFL); Upright Familiar Digit in Upright Familiar Letters, (UFD-in-UFL); Inverted Familiar Letter in Upright Familiar Letters (IFL-in-UFL); Upright Familiar Letter in Upright Non-familiar Letters (UFL-in-UNL); Upright Familiar Letter in Upright Familiar Digit (UFL-in-UFD); Upright Familiar Letter in Inverted Familiar Letters (UFL-in-IFL). The panels show discrimination responses: Z-scored summed spectra in source space masked by significant clusters from a cluster-based permutation test against zero.

Laterality indices were analyzed across broadly defined cortical regions (defined by the corresponding parcels from Desikan-Killiany atlas (‘aparc’), embedded in MNE-python, see Methods section for details), that included fusiform gyrus, inferior parietal region, inferior temporal lobule, lateral occipital cortex, lingual gyrus, middle temporal, parahippocampal gyri, superior temporal sulcus (‘banksst’) and primary auditory cortex region (‘transversetemporal’). Results at Fig. 4 show that for the majority of ROI, including fusiform gyrus, inferior and middle temporal regions, the discrimination response is significantly right lateralized. For the fusiform and inferior parietal area, left lateralized discrimination responses were observed in UFL-in-UFD categorical condition.

**Table 1.** Distribution of the significant discrimination response across brain regions in different experimental conditions. Asterisk sign “*” in a red cell indicates that the significant response was observed within the specified region in the given condition and empty blue cells indicate no significant response.

| Brain region | Left hemisphere |  |  |  |  |  |
| --- | --- | --- | --- | --- | --- | --- |
|  | UNL-in-UFL | UFD-in-UFL | IFL-in-UFL | UFL-in-UNL | UFL-in-UFD | UFL-in-IFL |
| Sup. temporal sulcus | * |  | * | * |  |  |
| Fusiform gyrus | * | * | * | * | * | * |
| Inf. parietal cortex | * | * | * | * | * | * |
| Inf. temporal cortex | * |  |  |  |  |  |
| Lat. occipital cortex | * | * | * | * | * | * |
| Lingual gyrus | * | * | * | * | * | * |
| Mid. temporal cortex | * |  |  |  |  |  |
| Parahippocampal gyrus | * |  |  | * | * |  |
| Pericalcarine gyrus | * | * | * | * | * | * |
| Post. cingulate cortex | * |  |  | * |  |  |
| Precuneus | * |  | * | * |  | * |
| Sup. parietal cortex |  |  |  | * |  |  |
| Primary auditory cortex | * |  |  |  |  |  |
| Brain region | Right hemisphere |  |  |  |  |  |
|  | UNL-in-UFL | UFD-in-UFL | IFL-in-UFL | UFL-in-UNL | UFL-in-UFD | UFL-in-IFL |
| Sup. temporal sulcus | * |  |  | * | * | * |
| Fusiform gyrus | * | * | * | * | * | * |
| Inf. parietal cortex | * | * | * | * | * | * |
| Inf. temporal cortex | * |  |  | * |  |  |
| Lat. occipital cortex | * | * | * | * | * | * |
| Lingual gyrus | * | * | * | * | * | * |
| Mid. temporal cortex | * |  |  | * |  |  |
| Parahippocampal gyrus | * |  |  | * | * |  |
| Pericalcarine gyrus | * | * | * | * | * | * |
| Post. cingulate cortex | * |  |  | * |  |  |
| Precuneus | * | * | * | * |  | * |
| Sup. parietal cortex |  |  |  | * |  |  |

\*Brain regions in accordance with labels of ‘aparc’ parcellation embedded in ‘mne’ python): ‘bankssts’ - banks of Superior Temporal Sulcus (STS), ‘fusiform’ - fusiform gyrus, ‘inferiorparietal’ - Inferior Parietal cortex, ‘inferiortemporal’ - Inferior temporal cortex, ‘lateraloccipital’ - Lateral Occipital cortex, ‘lingual’ - Lingual gyrus, ‘middletemporal’ - Middle temporal cortex, ‘parahippocampal’ - Parahippocampal gyrus, ‘pericalcarine’ - Pericalcarine gyrus, ‘posterior cingulate’ - Posterior Cingulate cortex, ‘precuneus’ - Precuneus, ‘superiorparietal’ - Superior parietal cortex, ‘transversetemporal’ - Primary auditory cortex.

**Fig. 4.**
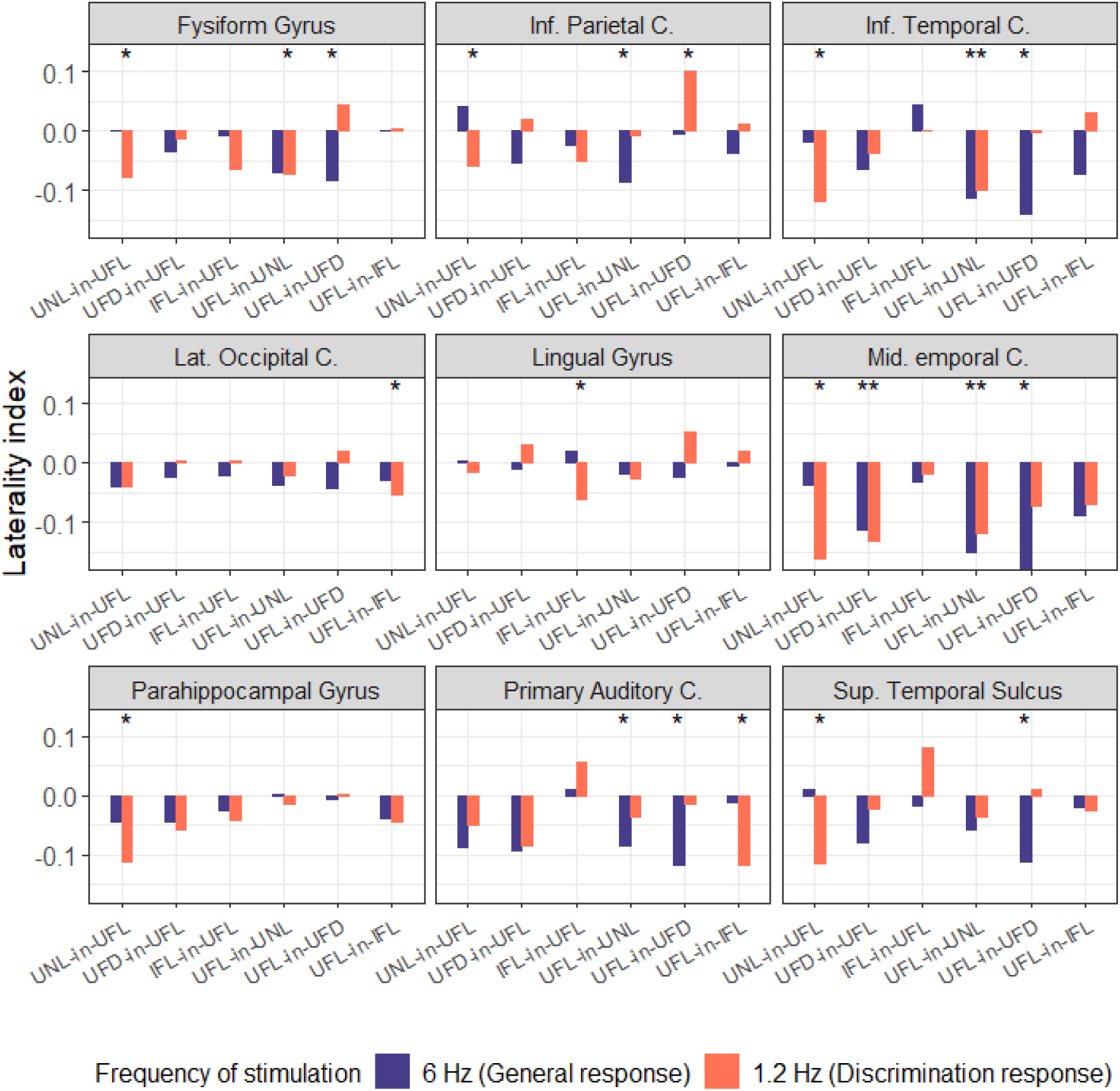
Mean laterality index (*LI* = (*LH* − *RH*)/(*LH* + *RH*)), calculated using Z values, for selected brain regions (“C. denotes “Cortex”), in different conditions and considering base (6 Hz) and deviant (1.2 Hz) stimulation frequency, significant responses marked with asterisks (*). LI value > 0 indicates left lateralization, while LI value < 0 indicates right lateralization.

### Category discrimination and reaction to novelty

To analyze the differences between novelty reaction and categorical discrimination, we compared the condition with upright non-familiar (foreign) letters (UNL-in-UFL, UFL-in-UNL) to the condition with upright familiar digits (UFD-in-UFL, UFL-in-UFD). Significant differences were observed for comparison of conditions where upright familiar letters were used as base category (UNL-in-UFL vs UFD-in-UFL), but not in the switched conditions, indicating better sensitivity to category vs novelty contrast of the conditions when different stimuli used as deviant. The major differences in localization for that comparison (UNL-in-UFL > UFD-in-UFL) include clusters in the right middle lateral occipitotemporal (BA39), inferior parietal (BA39) cortices, posterior supramarginal (BA39), lingual (BA18) and fusiform (BA37) gyrus (see Fig. 5), indicating that these regions are more specifically tuned to familiarity but not category of stimulus. The details of the found clusters of differences in the localization discrimination responses are summed up in Table S1 in the Supplementary materials.

**Fig. 5.**
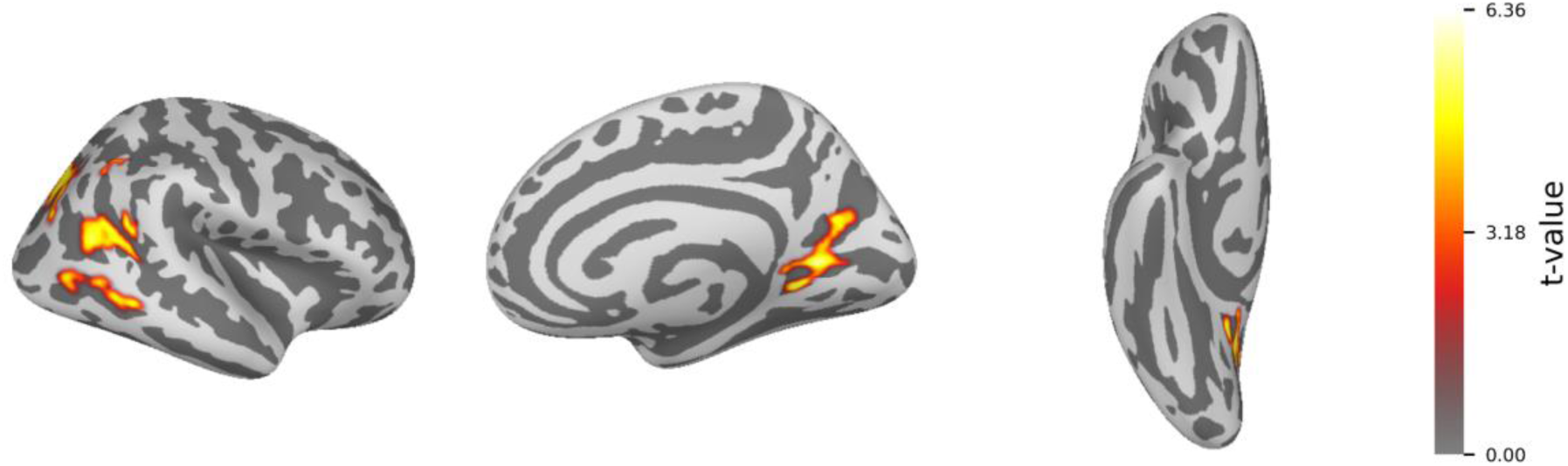
Significant clusters that differ the UNL-in-UFL (novelty) vs UFD-in-UFL (category) condition (UNL-in-UFL > UFD-in-UFL) were found only for the right hemisphere, t-values.

### Orientation discrimination and reaction to novelty

To search for specific brain regions responsible for orientation tuning we explore two contrasts. At first, we present data on orientation vs novelty (IFL-in-UFL vs UNL-in-UFL, UFL-in-IFL vs UFL-in-UNL).

Comparison UNL-in-UFL vs IFL-in-UFL (UNL-in-UFL > IFL-in-UFL) demonstrate significant clusters that include the ventral occipitotemporal regions and bilateral fusiform area, indicating its specific involvement in familiarity/novelty detection, but not orientation tuning (Fig. 6). Overall, this contrast indicates largely overlapping brain systems subserving these processes with orientation tuning being a subsystem of a more widely spread familiarity/novelty detection system.

**Fig. 6.**
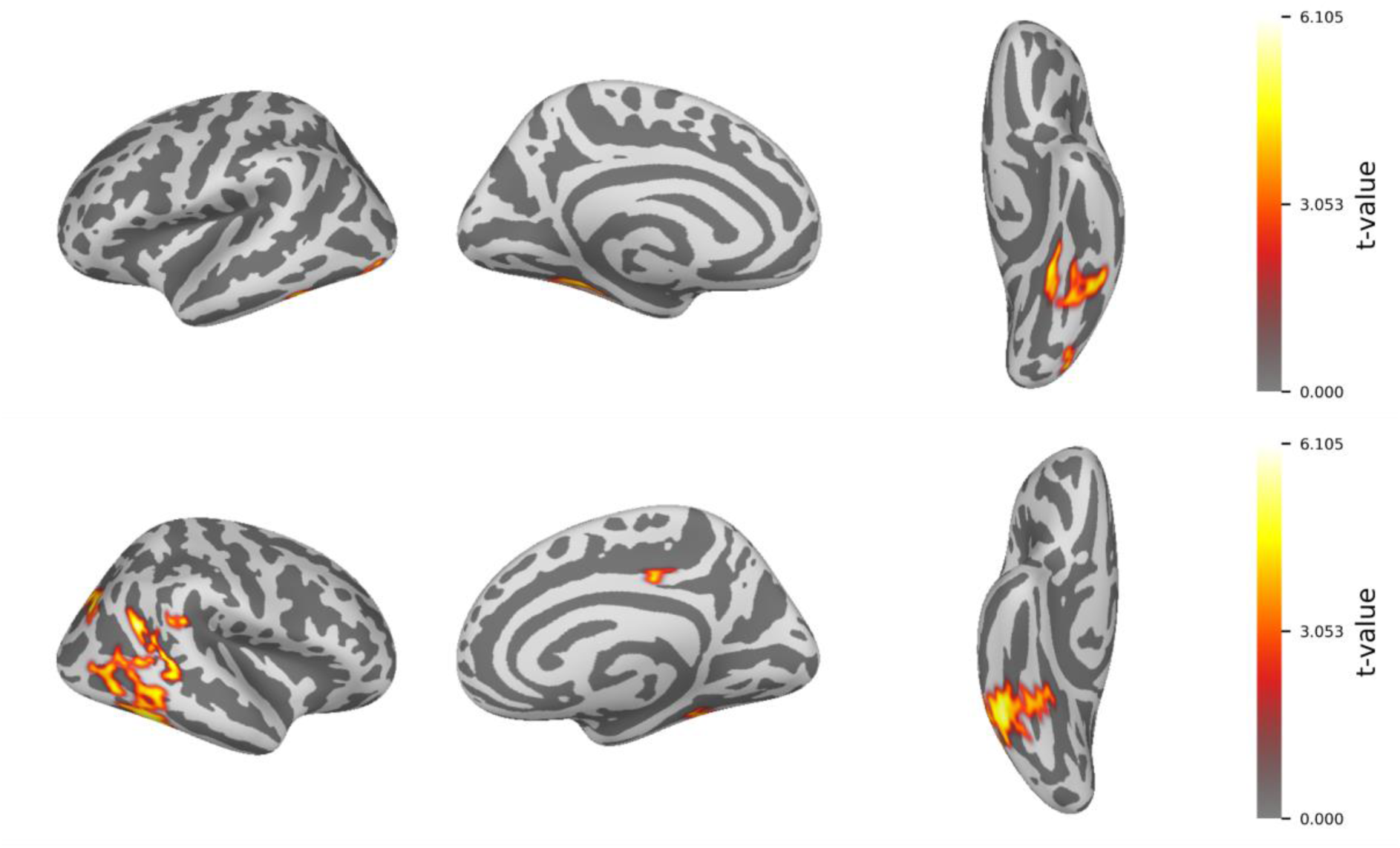
Significant clusters that differ the UNL-in-UFL from IFL-in-UFL condition (UNL-in-UFL > IFL-in-UFL), t-values.

Contrasting UFL-in-UNL vs UFL-in-IFL revealed significant clusters both in the left and the right hemispheres (Fig. 7), the spatial features of which have much in common with the UNL-in-UFL vs IFL-in-UFL contrast, including clusters in the right ventral temporo-occipital part and right fusiform gyrus (see Table S1 in Supplementary materials for description of the clusters and their characteristics).

**Fig. 7.**
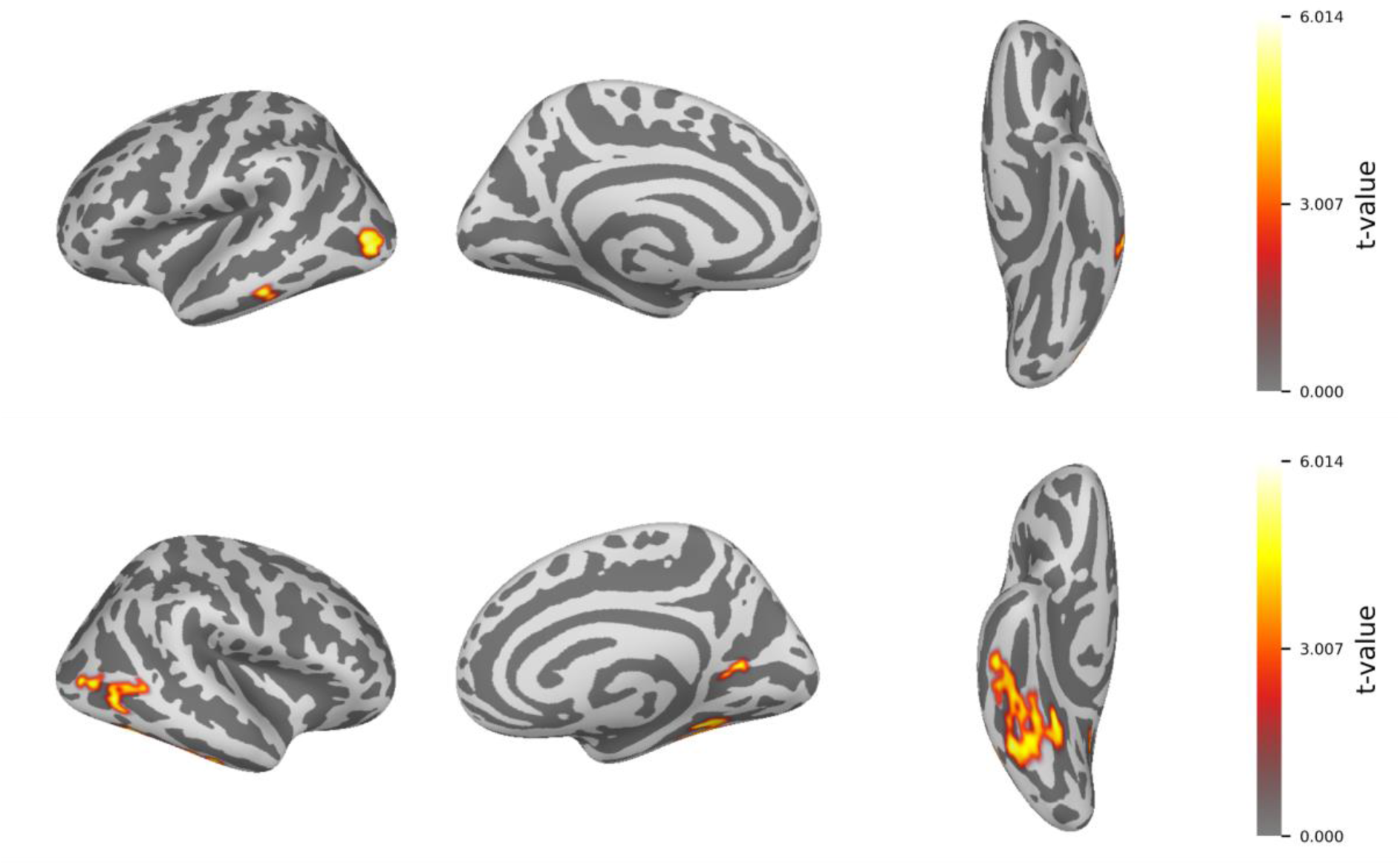
Significant clusters for comparison of UFL-in-UNL vs UFL-in-IFL conditions (UFL-in-UNL > UFL-in-IFL), t-values.

### Orientation and category discrimination

For orientation vs category tuning, no significant clusters were found in the comparison of conditions with upright familiar letters as standard stimuli (UFD-in-UFL vs IFL-in-UFL). The comparison between conditions using upright familiar letters used as deviant stimuli (UFL-in-UFD vs UFL-in-IFL) showed the cluster in the left lateral occipital cortex (BA 18/19), that was more active in UFL-in-UFD category condition, but was not activated in UFL-in-IFL novelty condition (Fig. 8).

**Fig. 8.**
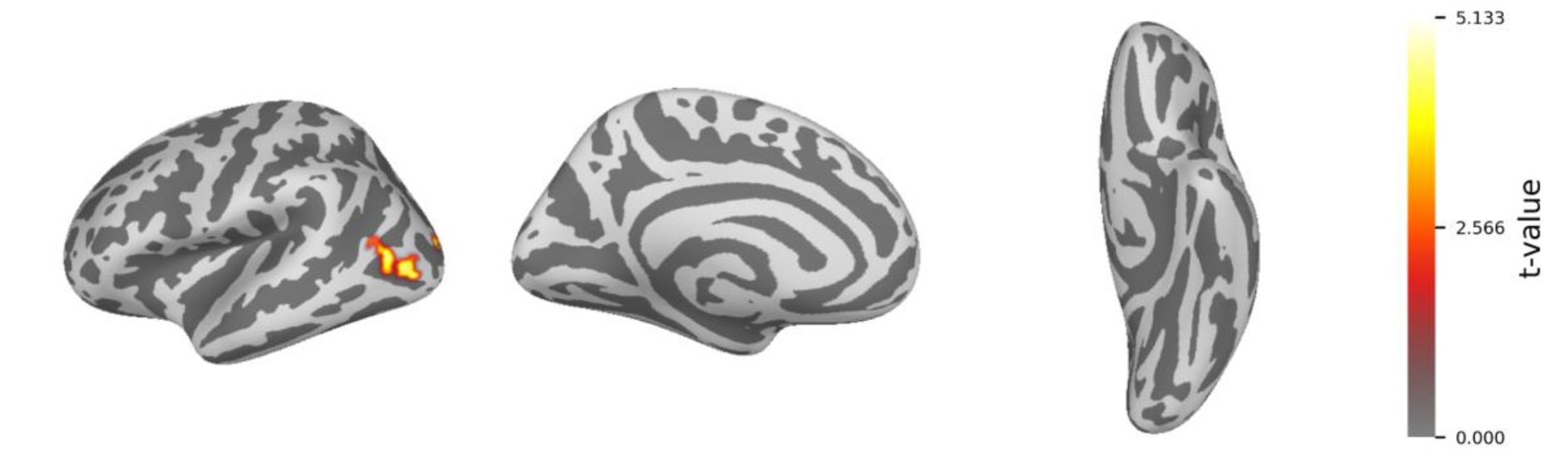
Significant clusters for comparison of UFL-in-UFD vs UFL-in-IFL conditions (UFL-in-UFD > UFL-in-IFL), t-values.

### Context related discrimination: contrasting switched blocks

Statistical comparison for switched pairs of conditions (UNL-in-UFL vs UFL-in-UNL, IFL-in-UFL vs UFL-in-IFL, UFD-in-UFL vs UFL-vs-UFD) was performed to estimate the role of standard and deviant category on discrimination response. Comparisons of UNL-in-UFL vs UFL-in-UNL conditions and IFL-in-UFL vs UFL-in-IFL conditions revealed no significant differences, suggesting pure discrimination response, which is independent of the context for novelty and orientation tuning. Comparison of UFD-in-UFL vs UFL-in-UFD category conditions (UFD-in-UFL < UFL-in-UFD) revealed 3 clusters in the left hemisphere, covering left fusiform, inferior temporal and lingual gyrus (Fig. 9), suggesting that differentiation of a category is context dependent and that these regions in the left hemisphere are more activated when familiar letters are presented in a stream of digits.

**Fig. 9.**
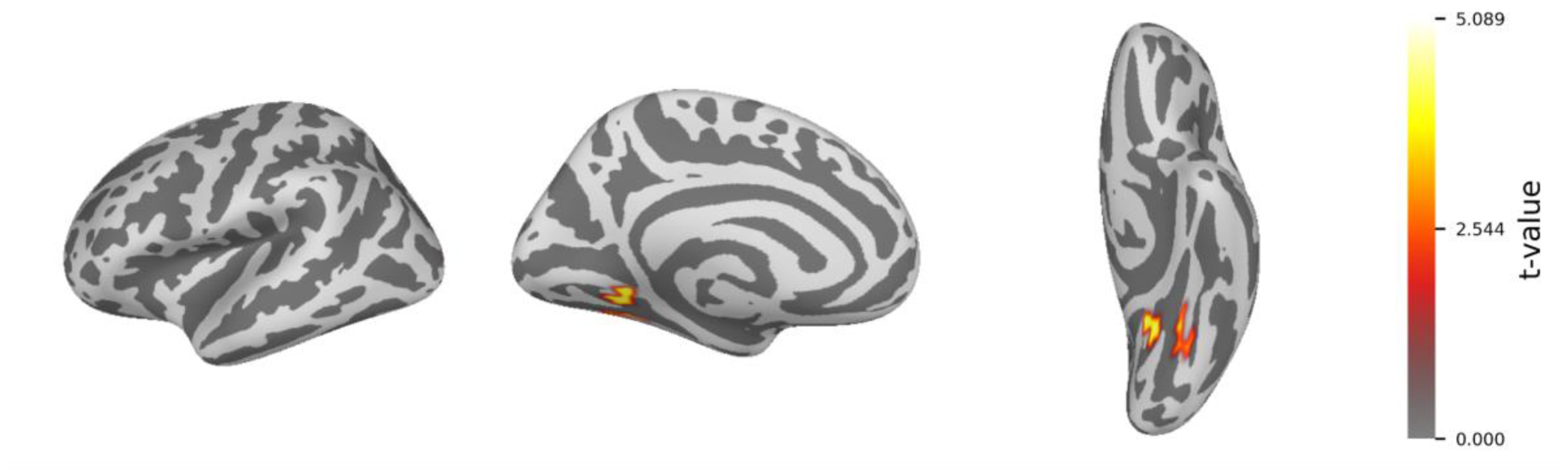
Significant clusters for comparison of UFL-in-UFD vs UFD-in-UFL conditions (UFL-in-UFD > UFD-in-UFL), t-values.

## Discussion

### “Embedded” organization of tuning for novelty and tuning for orientation system

Aligned with previous work, we observed the consistent discrimination response to novelty (UNL-in-UFL and UFL-in-UNL conditions) (De Rosa et al., 2024; Hauk et al., 2025; Lutz et al., 2024; Van De Walle De Ghelcke et al., 2021), and category (UFD-in-UFL and UFL-in-UFD conditions) (Lochy & Schiltz, 2019), and less pronounced tuning for orientation responses in conditions with inverted native letters (IFL-in-UFL and UFL-in-IFL), see Fig. 3. The observed discrimination responses showed gradual increase from orientation tuning, followed by category tuning with maximum and more distributed response for novelty tuning (UNL-in-UFL and UFL-in-UNL > UFD-in-UFL and UFL-in-UFD > IFL-in-UFL and UFL-in-IFL). Such increase in discrimination response between conditions is consistent with findings where more ‘distant’ (or contrastive) stimuli elicited higher and more distributed response and was observed for different stimuli. In particular, higher in amplitude neuronal response were registered for more higher-level categories, such as live vs non-live compared to animals vs plants in conceptual categorization (Stothart et al., 2017), larger difference in number magnitude (Marlair et al., 2022) and for strings of pseudoletters within words if compared with strings of consonants within words (Lutz et al., 2024). Thus, we may suggest that discrimination of familiar vs non-familiar letter is more salient in terms of brain activity, than between category differentiation (letters vs digits), with the smallest discrimination response for orientation discrimination.

It is still unclear whether higher response amplitudes may reflect local dynamics and integration between regions (Tewarie et al., 2019). However, the observed topography and the condition-specific response differences suggest a distributed organization with specialized modules for specific functions, such as novelty response, categorical differentiation, tuning for letters, tuning for orientation. It is supposed that the broader recruitment of neural systems during processing of unfamiliar and therefore more salient stimuli is associated with higher attention demands (Lochy et al., 2024). In contrast, more specialized systems are required for stimuli with which the subject is already familiar. For instance, foreign-script letters may activate the ventral visual stream and language-specific regions involved in processing potential linguistic content, analogous to how reading expertise refines the N170 component topography (Maurer et al., 2005). Inverted letters may engage narrower, more specialized and later developed neuronal system, related to specific expertise with particular stimuli (letters) and grounded within the already specialized letter recognition system. In line with the neuronal recycling theory (Dehaene & Cohen, 2007), which posits that novel functions may utilize and ‘recycle’ existing brain circuitry, we propose that unfamiliar linguistic stimuli engage the same structures as familiar characters, but with greater attentional and visual processing demands, that potentially explains the “embedded” topography of responses observed in our conditions.

### Lateralization of single letters processing is not limited to the left hemisphere

Our results show the mixed lateralization for different conditions with the presence of significant right lateralized discrimination responses in UNL-in-UFL and UFL-in-UNL novelty conditions for the majority of regions of interest (Fig. 3). The left lateralized responses were observed only in inferior parietal cortex for UFL-in-UFD category condition. While studies using the FPVS approach mostly showed left lateralized discrimination response in sensor space (Lochy et al., 2015, 2016; Rossion & Lochy, 2022; Volfart et al., 2021; Wang et al., 2021, 2023), source space analysis demonstrates bilateral discrimination response for words in consonant strings or non-words (Hauk et al., 2025) or for words in pseudofonts (De Rosa et al., 2024), which is consistent with our findings.

The specific mechanisms of the right hemisphere engagement in visual language processing remain unclear (Rossion & Lochy, 2022). Bilateral response may be suggested to be the part of genuine orthographic processing (see, for example, (Chu & Meltzer, 2019; Cohen et al., 2003)). According to Cohen’s visual word processing model (Cohen et al., 2003), letters are first analyzed involving information processing in occipitotemporal areas of both hemispheres, where increasingly abstract representations are computed. However, the left and right VWFA may serve different functional roles. The left VWFA is thought to process the letter identities, invariant to specific case and font representation, while the right VWFA may contribute to letter by letter processing in reading (for example in dyslexia (Cohen et al., 2003)), or be modulated by spatial attention. The right VWFA could specialize in processing character orientation included in its spatial and shape configuration, conjoining multiple visual features (Dien, 2009). This model is supported by findings on the interhemispheric connectivity during word recognition, which involves multiple processing stages. Specifically, directed connections from the right to left middle occipital gyri and reciprocal bidirectional connections between the left and right visual word form areas (Chu & Meltzer, 2019) provide evidence for right hemisphere involvement in early stages of lexical processing.

There is also evidence for ambiguous and changing hemispheric lateralization of language processing during development (Amora et al., 2022; Lutz et al., 2024; Olulade et al., 2020). In previous work (Kostanian et al., 2025), right-sided lateralisation of the discrimination response for identical novelty contrast (non-native non-familiar Georgian letters within native language Russian letters) was observed in children aged 3 to 9 years. One possible explanation for that result could be related to the young age of the participants, as brain responses to letter stimuli in children may be bilateral (Eberhard-Moscicka et al., 2015) or right-sided (Maurer et al., 2005), due to a not yet fully formed VWFA (Maurer & McCandliss, 2007). The confirmation of these results in adults rejects this hypothesis and requires alternative explanations for this phenomenon.

While the overall response to deviant stimuli was predominantly bilateral or had a tendency to be right lateralized in specific brain areas, cross-block comparisons allow us to identify the precise spatial and magnitude differences in neural tuning in the conditions which corresponded with novel stimuli discrimination, categorical detection or tuning to the certain orientation. The results suggest that the right lateralized response may be primarily attributed to the novelty reaction, while discrimination of native letters in a stream of digits as compared to switched condition were supported by functionally specialized left hemisphere structures particularly the ventral occipitotemporal cortex (vOTC) and lingual gyrus. The results are summarized in the scheme below (Fig. 10).

**Fig. 10.**
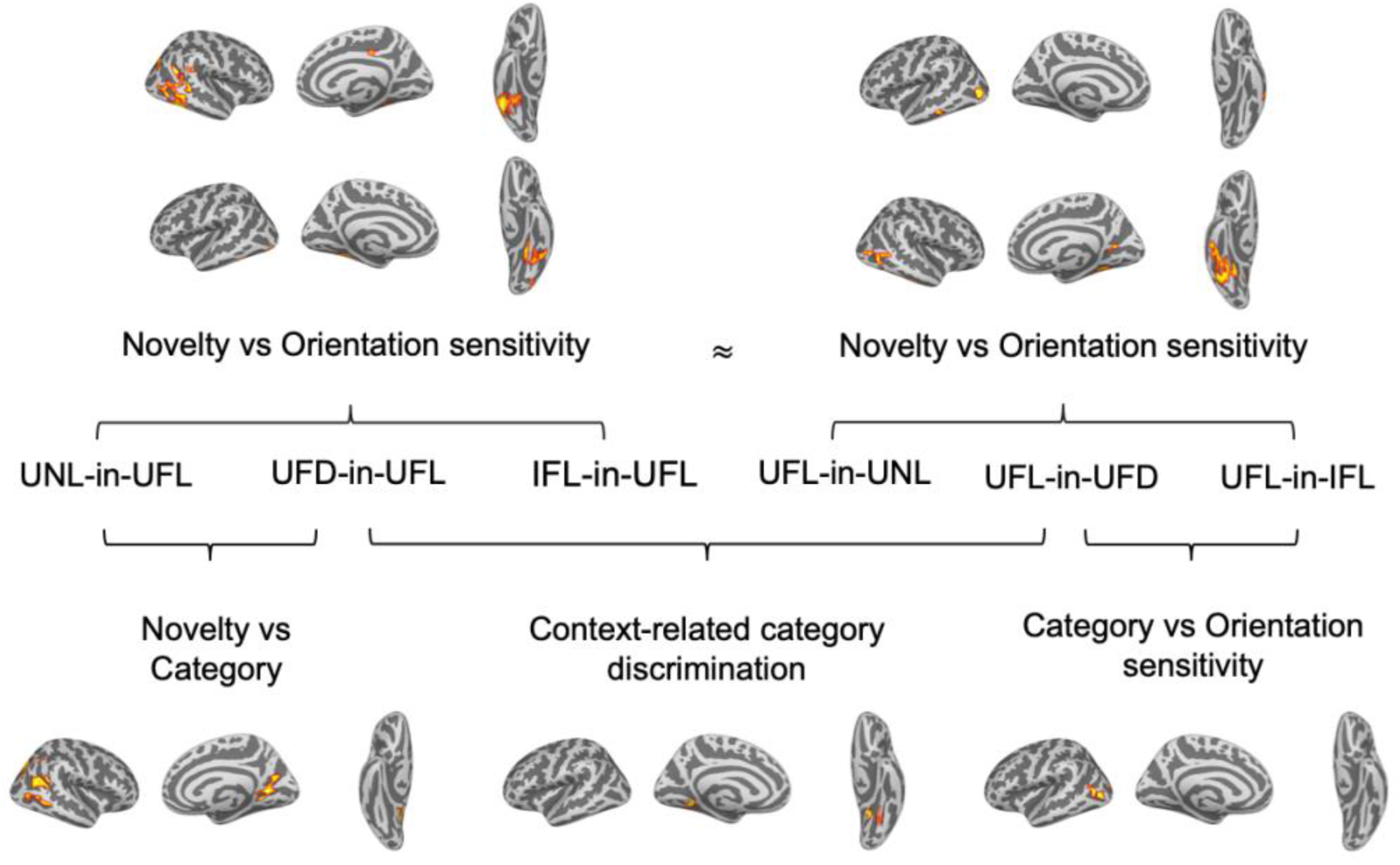
The scheme of contrasts and interpretation for pairwise comparisons.

### Familiarity vs category tuning

According to our hypothesis, comparing UNL-in-UFL and UFD-in-UFL conditions could help us reveal the specific structures associated with novelty and category discrimination. While both digits and Georgian letters are not letters of native alphabet, digits in contrast to the letters of the Georgian alphabet, are familiar and well-known characters to adult participants. Significant clusters revealed in novelty (UNL-in-UFL) vs category (UFD-in-UFL) conditions comparison (UNL-in-UFL > UFD-in-UFL) include the area in the right inferior temporo-parietal region, comprising temporoparietal junction (TPJ) and angular gyrus (AG). According to the literature, similar activation of the right TPJ/AG is also observed as a reaction to novelty, for example when a salient contextual cue facilitates target discrimination (Geng & Mangun, 2011), in case of reorienting of attention (Krall et al., 2015) and when infrequent deviant stimuli are presented within a series of standard events (Vossel et al., 2009), supporting visual attention processes involvement (Devaney et al., 2019). Greater right than left-sited activations in temporal and parietal regions was observed for target detection and novelty processing during auditory oddball paradigm (Stevens et al., 2005). The involvement of right AG can also reflect the engagement of attentional control as part of the implementation of executive functions along with domain specific language processing (Kim et al., 2022).

Therefore, it can be assumed that the stage of coarse tuning for print, which is often associated with categorical differentiation, may be preceded by a level of prime tuning for familiarity. This level is not associated with specific stimuli categories and is established through perceptual experience and interaction with various stimuli in the environment. Thus, the discrimination response to previously unfamiliar non-letter stimuli (e.g., foreign language letters or pseudofont characters) may represent different aspects of character processing, including category discrimination for language and non-language stimuli itself and response to novelty, which is important to keep in mind using such stimuli categories as oddball stimuli in FPVS studies.

### Tuning for letter orientation in adults

It is generally accepted that developing reading proficiency requires the ability to form representations of letters in their specific orientations. Thus, the invariance that may be beneficial for object recognition must be overcome during reading acquisition to enable more effective orthographic characters recognition (Dehaene et al., 2010; Pegado et al., 2011). Previous studies showed that the visual system establishes specialized mechanisms for letter orientation discrimination (Pegado et al., 2011), that may differ in letter or word recognition but share some common principles including spatial attention, supporting invariance (Rossion & Lochy, 2022) and require a sufficient level of expertise (Kostanian et al., 2025). We attribute this sensitivity to special tuning for letters orientation. So, reading expertise is characterized by the development of a specialized system that actively suppresses general invariance mechanisms to become sensitive to the specific orientations that are critical for efficient orthographic processing.

In the present study, we were able to observe a discrimination response for inverted native language letters among upright letters, as well as in the opposite condition for upright letters among inverted letters in adults (Fig. 2). However, comparison with other experimental conditions did not allow us to identify specific structures that could be devoted solely to the inverted letter perception. That may be explained by the overlap of the neural basis for coarse and orientation tuning, with the former involving specialized mechanisms for sensitivity to letters orientation. For instance, it was shown that VWFA, known for category-specific word/letter response, also plays a crucial role in mirror-invariance resolution for letters. Pegado et al. (2011) using fMRI and priming paradigm showed that VWFA is critical in distinguishing the left-right orientation of single letters in skilled readers, and yet exhibits mirror invariance for simple pictures of equivalent complexity. The work of (Nakamura et al., 2014) revealed that temporarily disrupting the left ventral occipitotemporal cortex (VOTC) significantly delayed mirror-image recognition for letter-strings, but not for other objects, supporting its causal role in mirror-invariance suppression that is specific to letters.

However, the difference in coarse and orientation tuning, though sharing common neural substrate, may be attributed to special spatiotemporal organization of the category-specific discrimination response, for example accounting for the processes of suppression mirror-invariance mechanisms, that yield in the lower amplitude response in the orientation conditions. One possible mechanism for this suggests that left VWFA may contain a local inhibitory circuit that suppresses mirror-image generalization specifically for letters and words (Nakamura et al., 2014). This could explain the delayed N170 latencies observed for mirrored versus canonical (non-mirrored) representation, a phenomenon similar to the face inversion effect (Núñez-Peña & Aznar-Casanova, 2009; Tanaka, 2020). The event related potentials (ERP) studies’ findings (Rossion et al., 2003) showed left-lateralized N170 inversion effect for words in lateral inferior occipital cortex/posterior fusiform gyrus both for scalp topographies and source analyses, similar to face-like N170 inversion effect.

Evidence from ERP EEG studies supports the developmental expertise-dependent nature of specialized orientation tuning for reading. For instance, adults, but not children, exhibit distinct neural activation patterns to letter orientation, including recruitment of left VWFA (Blackburne et al., 2014). This suggests a transition from a broad, coarse tuning system to a specialized one that overcomes perceptual invariance. Furthermore, this expertise generalizes as literate individuals show increased orientation sensitivity to other visual categories like faces (Rossion & Lochy, 2022).

### Role of base and deviant category on discrimination response generation

Previous FPVS studies rarely used switched conditions, where the same stimulus categories were used as both standards and deviants. In the study by Lutz and colleagues (Lutz et al., 2024), the use of the words as deviant and base stimuli allowed to introduce the concepts of familiarity (real words, pseudowords or strings of letters as a deviant among pseudofont strings) and sensitivity (pseudofont strings as a deviant among real words). While the sensitivity model reflected general discrimination associated with the coarse tuning, the familiarity model allowed to test discrimination of more familiar oddball stimulus inside other letter strings. Familiarity model (familiar oddball stimulus e.g. words, pseudowords or consonant strings) provides more significant discrimination response on more number of harmonics, than sensitivity model (where unfamiliar oddball stimuli are used as base). This distinction raises the question about what process reflects a discrimination response: identification of a specific stimulus category or reaction to difference between two stimuli categories?

In our work the comparison of switched blocks aimed to help us answer the question of whether the category of stimulus used as standard or deviant stimuli impacts the features of the discrimination response. For most conditions the exchange between deviant and standard category did not influence the pattern of brain activation. The only difference we get in analysis of switched conditions is the left hemispheric cluster including left fusiform, inferior temporal and lingual gyrus for UFL-in-UFD vs UFD-in-UFL category contrast (UFL-in-UFD > UFD-in-UFL). The observed leftward lateralization of the discrimination response to letters among digits, compared to the discrimination response to digits among letters, support the first hypothesis (sensitivity to a certain category of stimuli). At the same time, the absence of differences between the remaining pairs of switched counterbalance conditions is more in favor of the second hypothesis (general reaction to differences). Assuming that difference between UFL-in-UFD and UFD-in-UFL category conditions may be associated differentiation between different categorical stimuli, we may also cautiously speculate that the absence of difference in other switched conditions with novelty and orientation conditions is associated with the similar cognitive and linguistic expectations, regardless of the origin or unusual orientation of a letter.

Another study of Lochy and Schiltz (2019) productively used the switched conditions to show the differences in lateralization of discrimination response for digits and letters. The distinct lateralization patterns observed for letters and digits are thought to reflect culturally acquired specialization for these characters, which tend to develop preferentially in different hemispheres (Lochy & Schiltz, 2019; Park et al., 2012). Our results are partly consistent with the evidence of mostly left lateralization for letters than for numbers discrimination (Lochy & Schiltz, 2019; Marlair et al., 2022; Park et al., 2012). However we did not find any significant clusters where activation for UFD-in-UFL would be more than for UFL-in-UFD, which could be due to the proposed bilateral processing of digits (Jung et al., 2020), that is also supported by contralateral connectivity evidence (Hannagan et al., 2015).

### Limitations and future directions

Several limitations of the current study pertain to the stimuli choice. It can be assumed that the topography of the discrimination response to non-letter stimuli may be influenced by both the degree of familiarity and the perceptual complexity of the stimuli. As novelty factors were established in familiarity contrast, category contrast (UFD-in-UFL, UFL-in-UFD) may be potentially influenced by level of visual processing, associated with different visual complexity. According to perimetric complexity estimates (see Methods), letters tend to be more complex visual patterns than digits (it was also shown that digits usually consist of fewer strokes than letters (Schubert, 2017)). These factors should be carefully considered when developing future stimulus materials. However, despite the fact that more complex stimuli, such as letters may yield a slightly higher SNR due to the greater number of visual features, we may expect that the localization of categorical response will remain. As it was shown in (Georges et al., 2020), similar electrophysiological response patterns were observed for numerical identities presented as either dots or pictures, regardless of stimulus complexity. Response observed in switched conditions with digits and upright native letters supports the view that the response to these stimuli reflects the neural signatures of distinct familiarized categories.

There is also a topic of interest associated with the neural mechanisms of the discrimination response and FPVS results interpretation, whether we can confidently attribute the response to deviant stimuli for the discriminative ability of certain categories of stimuli, and to what extent the implicit learning of statistical regularities in visual input streams may play a role (De Rosa et al., 2022). The latter interpretation may be particularly relevant to our study, as we used a small set of stimuli (N=8) for each category. It was shown that statistical regularities, associated with the different design-related factors (as set size and item repetition regularities), do not affect the robustness of the FPVS response (Lochy et al., 2024), however the impact of the context-dependent neural adaptation regarding different stimuli types and their relative positioning may be a question for the future research.

## Conclusion

Using the FPVS approach, we investigated the differences in familiarization (native vs non-native letters), category (letters vs digits) and orientation (upright vs inverted) tuning to letters. The topography of the responses showed a predominantly bilateral organization, encompassing regions specifically associated with the identification and processing of speech-related stimuli (e.g. left VOTC, STS, middle temporal areas). The overall pattern of results suggested embedded organization of different levels of visual stimuli analysis, with more broader areas activated for novelty detection, followed by that of category discrimination with the more local activation for orientation tuning.

Comparisons between novelty and categorical conditions revealed clusters in the right inferior temporo-parietal region, including the temporoparietal junction and angular gyrus, which may underlie neural responses to novelty. Contrasts between switched categorical conditions (letters vs digits) indicated a left-lateralized occipital cortex discrimination response to letters among digits, supporting specialization of this area for letter processing. However, no significant differences emerged between other switched condition pairs, suggesting context-independent coding for novelty and orientation. Our analysis also did not reveal the specific spatial patterns associated with orientation tuning, suggesting that orientation-specific tuning mechanisms may operate within an embedded framework — spatially overlapping with coarse tuning networks of category detection but exhibiting unique spatiotemporal dynamics. Further research is required to elucidate these fine-grained tuning mechanisms and their functional organization within the visual processing hierarchy.

## Acknowledgements

We are especially grateful to Andrey Prokofyev, Anastasia Neklyudova, Nikita Korotkov, Diana Smolskaya, Ekaterina Slovenko for help with MEG data collection, Gurgen Soghoyan for advisement in data analysis, and we also wish to thank all participants for their contribution to this study.

## Declarations

Funding: This work was supported by Russian Science Foundation (RSF), grant №23-78-00011. Competing interests: The authors report no competing interests to declare.

Consent: All participants have given written informed consent.

Ethics approval: The study, approved by the Institute of Higher Nervous Activity and Neurophysiology’s ethics committee (June 30, 2023), followed the Helsinki Declaration’s ethical guidelines for human research. Data/Code Availability: The data of this study is available from the corresponding author upon reasonable request. The code for the study analysis is available at https://github.com/katerkoch/fpvs_meg_letters

