## Supplementary materials for "Novelty, category and orientation tuning for printed characters: a magnetoencephalography study with fast periodic visual stimulation"

Journal name: Brain Topography

Kochetkova, E.*, Kostanian, D., Martynova, O., Sysoeva, O.
* Faculty of Biology and Biotechnology, HSE University; Institute of Higher Nervous Activity and Neurophysiology RAS, Moscow, Russia;

**Supplementary materials**

**Table S1**: Significant clusters (p < 0.001 to form cluster, p < 0.05 for cluster in permutation t-test against zero) in source space. Notation «Condition1 > Condition2» stands for the higher activation in Condition1.

| Conditions | MNI coordinates of cluster's centroid | Number of vertices in a cluster | Mean cluster t-value (cluster p-value) | Hemisphere | Brain regions according to Harward-Oxford atlas with corresponding Brodmann area |
|---|---|---|---|---|---|
| UNL-in-UFL > UFD-in-UFL | 34, -71, 29 | 88 | 4.47 (0.001) | right | Lateral occipital cortex, middle (BA39) |
| | 43, -54, 42 | 16 | 4.12 (0.032) | right | Inferior parietal cortex (BA39) |
| | 25, -62, 1 | 86 | 4.5 (0.001) | right | Lingual gyrus (BA18) |
| | 45, -60, 21 | 71 | 4.49 (0.001) | right | Lateral occipital cortex, middle (BA39) |
| | 55, -58, 2 | 24 | 4.31 (0.015) | right | Middle Temporal Gyrus, Fusiform cortex (BA37) |
| | 46, -60, 2 | 14 | 4.2 (0.041) | right | Middle Temporal Gyrus, Fusiform cortex (BA37) |
| | 50, -44, 18 | 15 | 4.51 (0.031) | right | Supramarginal Gyrus, posterior division (BA39) |
| UNL-in-UFL > IFL-in-UFL | -42, -82, -13 | 14 | 4.1 (0.039) | left | Lateral Occipital Cortex, inferior (BA19) |
| | -36, -44, -21 | 39 | 4.08 (0.009) | left | Fusiform cortex (BA37) |
| | -32, -53, -10 | 28 | 4.16 (0.013) | left | Fusiform cortex (BA37) |
| | 46, -44, 8 | 51 | 4.32 (0.005) | right | Middle Temporal Gyrus, temporooccipital part (BA21) |
| | 34, -71, 29 | 53 | 4.24 (0.005) | right | Lateral occipital cortex, middle (BA39) |
| | 41, -48, -19 | 128 | 4.18 (0.001) | right | Fusiform cortex (BA37) |
| | 7, -27, 40 | 12 | 4.17 (0.048) | right | Cingulate Gyrus, posterior (BA31) |
| | 46, -56, 21 | 27 | 4.45 (0.013) | right | Angular Gyrus (BA39) |
| | 61, -37, 18 | 16 | 4.24 (0.029) | right | Supramarginal Gyrus, posterior (BA22) |

| Conditions | MNI coordinates of cluster’s centroid | Number of vertices in a cluster | Mean cluster t-value (cluster p-value) | Hemisphere | Brain regions according to Harward-Oxford atlas with corresponding Brodmann area |
|---|---|---|---|---|---|
| UFL-in-UNL > UFL-in-IFL | -43, -80, 0 | 38 | 4.6 (0.003) | left | Lateral Occipital Cortex, inferior (BA19) |
| | -57, -34, -16 | 12 | 4.49 (0.044) | left | Middle Temporal Gyrus, posterior (BA21) |
| | 44, -68, 7 | 15 | 4.47 (0.028) | right | Lateral Occipital Cortex, inferior (BA19) |
| | 41, -48, -19 | 129 | 4.27 (0.0001) | right | Fusiform cortex (BA37) |
| | 55, -58, 2 | 34 | 145.9 (0.005) | right | Fusiform cortex (BA37) |
| | 27, -65, 7 | 15 | 4.32 (0.031) | right | Intracalcarine Cortex (BA23) |
| UFL-in-UFD > UFL-in-IFL | -40, -68, 0 | 19 | 4.38 (0.02) | left | Lateral Occipital Cortex, inferior (BA19) |
| | -45, -63, 11 | 19 | 4.24 (0.018) | left | Middle Temporal Gyrus, temporooccipital (BA39) |
| | -30, -92, 10 | 18 | 4.18 (0.02) | left | Occipital pole (BA18) |
| UFL-in-UFD > UFD-in-UFL | -13, -97, 14 | 15 | 4.26 (0.026) | left | Occipital pole (BA18) |
| | -17, -47, -7 | 24 | 4.26 (0.008) | left | Lingual Gyrus (BA19) |
| | -30, -63, -14 | 13 | 4.12 (0.038) | left | Fusiform gyrus (BA37) |

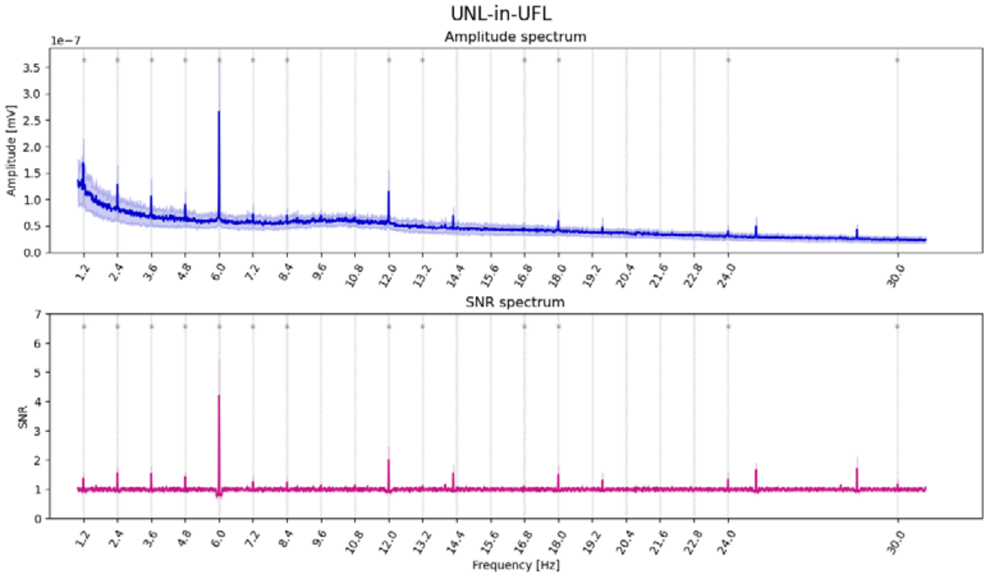

a) UNL-in-UFL

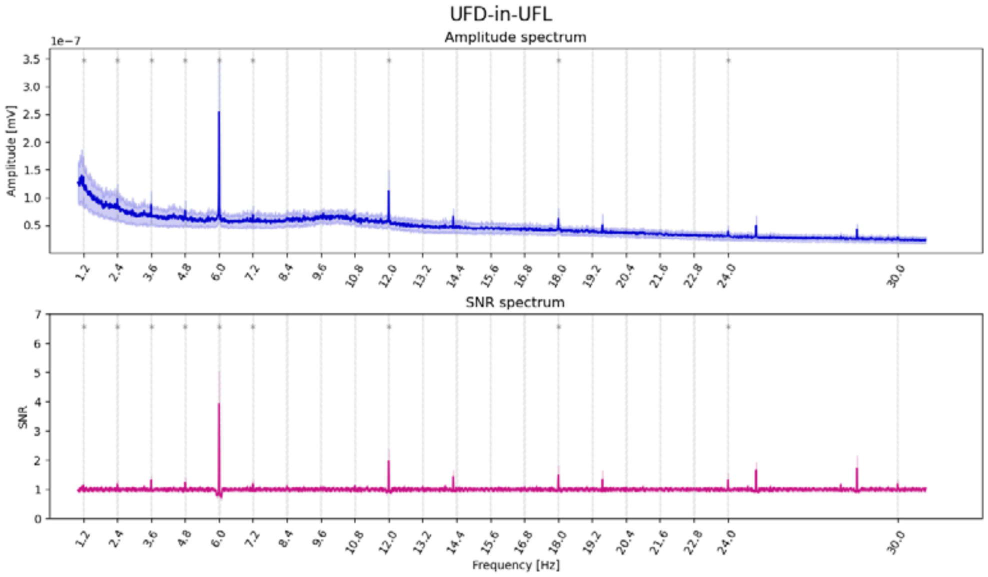

**b)** UFD-in-UFL condition

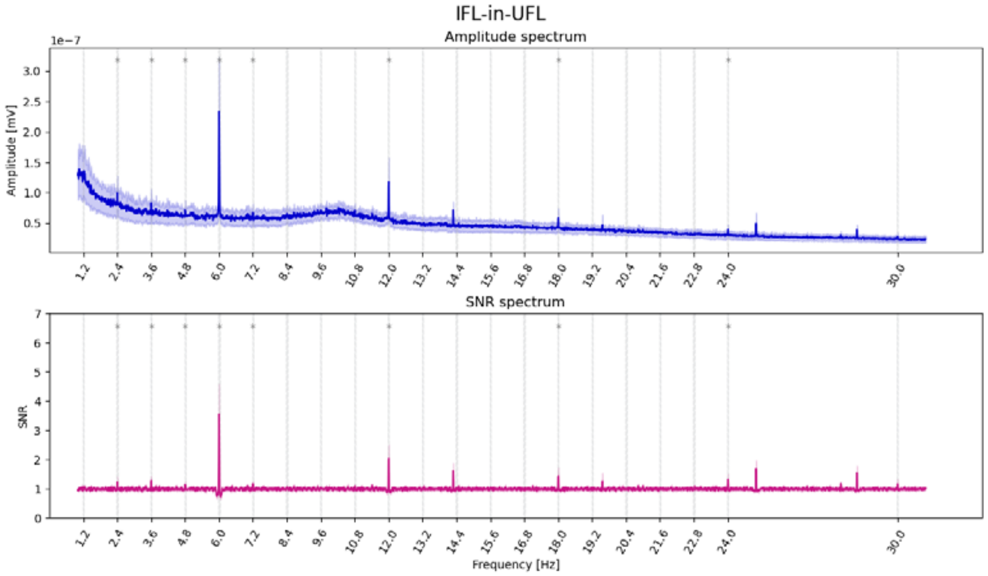


**c)** IFL-in-UFL condition

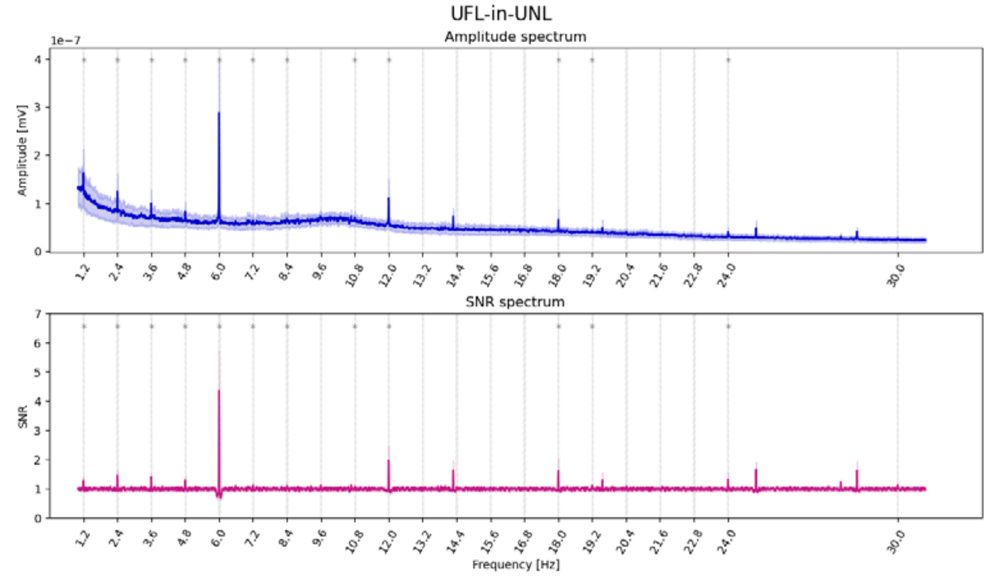


d) UFL-in-UNL

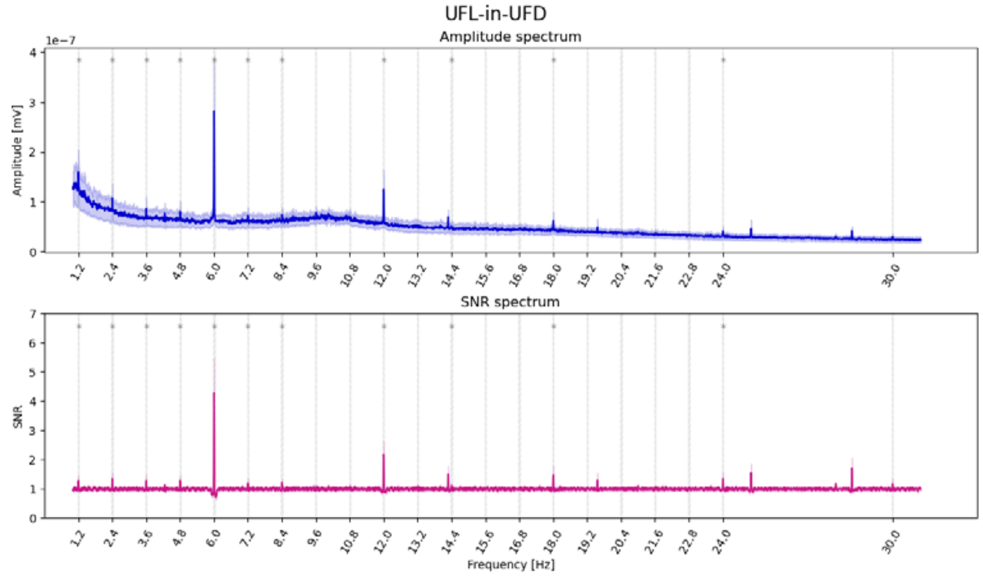


e) UFL-in-UFD

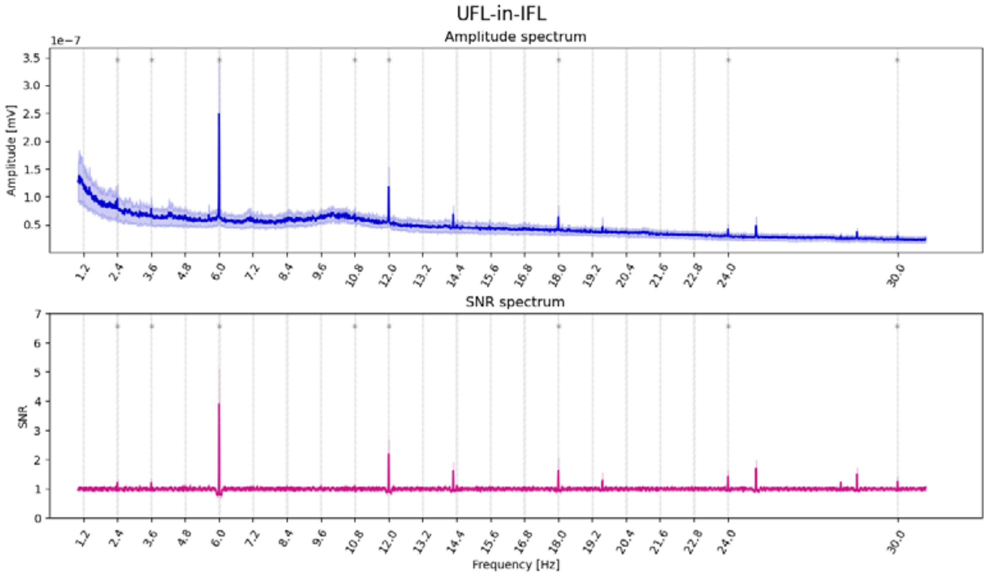


f) UFL-in-IFL

**Fig. S1** Amplitude and SNR spectrum for selected ROI (according to Desikan-Killiany atlas embedded in MNE parcellation: ‘lateraloccipital’, ‘fusiform’, ‘inferiorparietal’, ‘inferiortemporal’, ‘middletemporal’, ‘parahippocampal’, ‘lingual’, ‘transversetemporal’, ‘banksst’, ‘precuneus’, ‘pericalcarine’, ‘posteriorcingulate’) for conditions: a) UNL-in-UFL; b) UFD-in-UFL; c) IFL-in-UFL; d) UFL-in-UNL; e) UFL-in-UFD; f) UFL-in-IFL. The graph shows response at oddball stimulation frequency 1.2 Hz and its harmonics (in particular, 1.2 Hz, 2·1.2Hz = 2.4Hz, 2·1.2Hz = 3.6Hz, and the following 4.8Hz, 7.2Hz, 8.4Hz, 9.6Hz, 10.8Hz, 13.2Hz, 14.4Hz etc., vertical grey lines denote harmonics) and base stimulation frequency (6 Hz) and its first harmonics 12 Hz, 18 Hz, 24 Hz, 30 Hz).

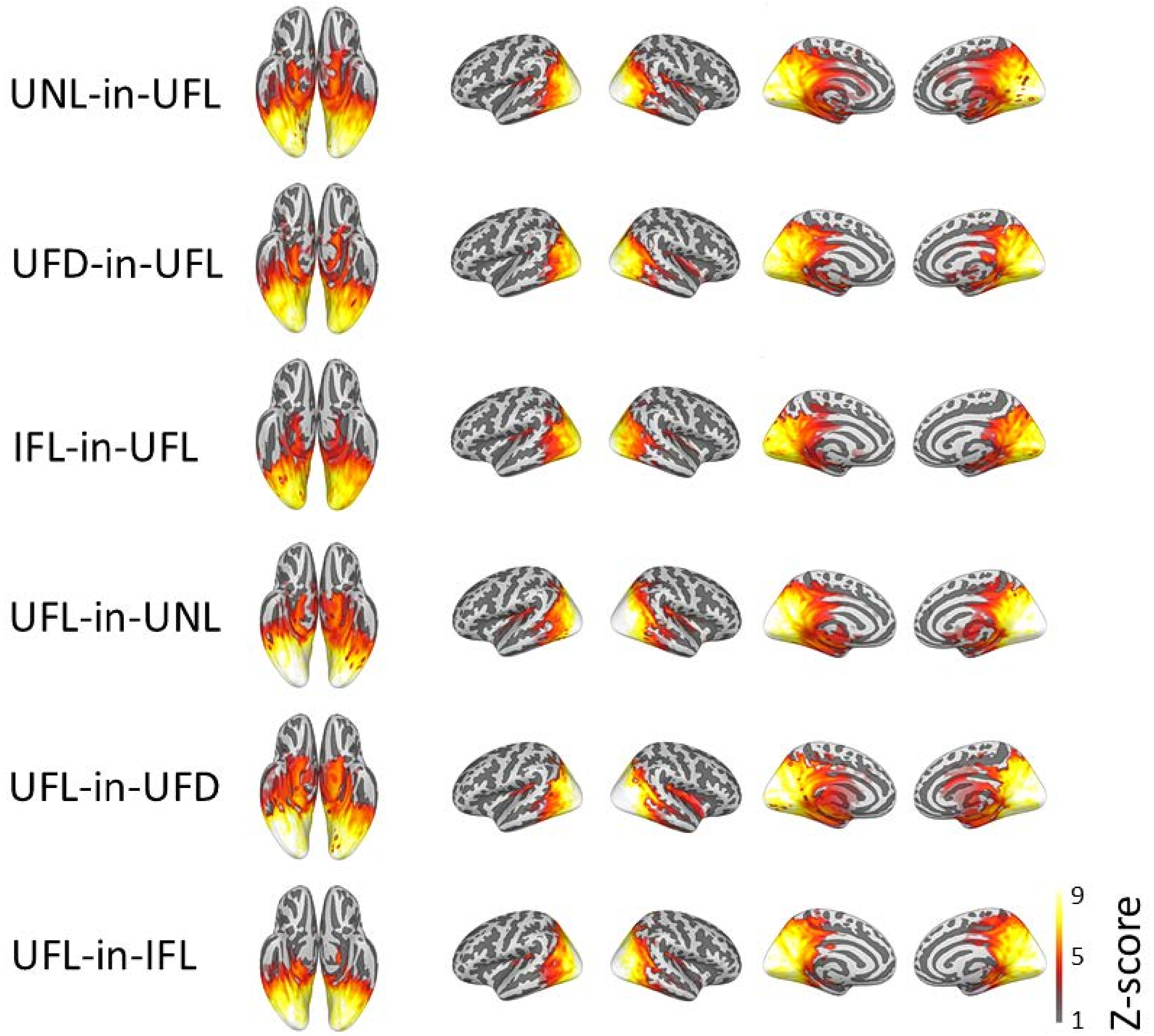


**Fig. S2** Z-scores for the summed base frequency (6 Hz) and its nine harmonics at all experimental conditions: Upright Non-familiar Letter in Upright Familiar Letters (UNL-in-UFL); Upright Familiar Digit in Upright Familiar Letters, (UFD-in-UFL); Inverted Familiar Letter in Upright Familiar Letters (IFL-in-UFL); Upright Familiar Letter in Upright Non-familiar Letters (UFL-in-UNL); Upright Familiar Letter in Upright Familiar Digit (UFL-in-UFD); Upright Familiar Letter in Inverted Familiar Letters (UFL-in-IFL). The panels show discrimination responses: Z-scored summed spectra in source space masked by significant clusters from a cluster-based permutation test against zero.
